# Blockade of Kv3.1 by MK-801, a PCP-Derivative NMDA Receptor Inhibitor: Implications for Models of Schizophrenia

**DOI:** 10.64898/2026.09.06.749750

**Authors:** Tae Jun Park, Sang Woong Park, Solah Park, Yeji Lim, Hyeryeong Lee, Jeongbin Kim, Harim Kim, Dong Hyun Kim, Jin O-Uchi, Mi Seon Seo, Jin Ryeol An, Bok Hee Choi, Young Min Bae

## Abstract

MK-801, a phencyclidine (PCP) derivative also known as dizocilpine, is a potent noncompetitive antagonist of the N-Methyl-D-aspartate receptor (NMDAr). The NMDAr plays a critical role in mediating excitatory synaptic transmission in the central nervous system (CNS) and is important in regulating synaptic plasticity, learning, and memory. MK-801 is known to induce schizophrenia-like phenotypes in animal models by blocking the NMDAr. Several studies have reported that the cognitive impairment associated with schizophrenia is linked to functional defects of ion channels in parvalbumin-positive GABAergic interneurons. Kv3.1 is a voltage-gated K^+^ (Kv) channel involved in the rapid repolarization of the action potential in neurons; it is richly expressed in parvalbumin-positive GABAergic interneurons and is associated with fast, repetitive spike generation. A decrease of Kv3.1 in the CNS has been reported to be associated with schizophrenia. In the present study, the effect of MK-801 on Kv3.1 was investigated using the whole-cell patch-clamp technique in Chinese hamster ovary (CHO) cells stably expressing Kv3.1. MK-801 caused a concentration-dependent inhibition of Kv3.1, with an IC50 of 10.81 μM and a Hill coefficient of 0.89. The blocking potency was stronger at depolarized potentials, indicating a voltage-dependent block. MK-801 also produced a use-dependent block, inducing progressive inhibition with repeated stimulation at increased frequencies (1 Hz and 2 Hz), consistent with a delay in recovery from inactivation of Kv3.1 in the presence of MK-801. In addition, MK-801 induced a hyperpolarizing shift in the voltage dependence of the steady-state inactivation curve of Kv3.1. Taken together, these results indicate that MK-801 blocks Kv3.1 expressed in CHO cells in a concentration-, voltage-, and state-dependent manner. Given the importance of Kv3.1 in parvalbumin-positive, fast-spiking GABAergic interneurons, these findings suggest that MK-801 may alter the firing patterns of these inhibitory neurons, contributing to the onset and symptoms of schizophrenia.

## Introduction

Schizophrenia is a chronic and debilitating psychiatric disorder characterized by a distorted perception of reality, encompassing positive symptoms (hallucinations, delusions), negative symptoms (social withdrawal, anhedonia), and cognitive deficits that collectively undermine daily functioning. The main causes are thought to involve an abnormal balance of neurotransmitters such as dopamine. In addition, genetic predisposition, abnormal neuroplasia, maternal infection during the fetal period, environmental and socio-cultural factors, stress, and trauma are also implicated. Most of these theories focus on excess or deficiency of neurotransmitters, including dopamine, serotonin, and glutamic acid [1–3]. Other theories implicate aspartic acid, glycine, and gamma-aminobutyric acid (GABA) as part of the neurochemical imbalance of schizophrenia [4,5]. One of the known contributors to schizophrenia pathogenesis is abnormal activity of the N-methyl-D-aspartate receptor (NMDAr), whose physiological agonist is glutamate [6,7]. Schizophrenia-like states can be induced by reduced glutamate/NMDAr function, and animal models induced by NMDAr blockers such as ketamine, MK-801, and PCP reportedly reproduce schizophrenia-like psychosis, including positive symptoms, negative symptoms, and cognitive dysfunction. Therefore, PCP- (or ketamine- and MK-801-) treated animals have been widely used to study the pathogenesis of schizophrenia [8–10].

PCP and its derivatives ketamine and MK-801 are structurally similar and classified as non-competitive NMDAr inhibitors. Among these, ketamine is used clinically as a dissociative anesthetic [11–14]. MK-801, which has effects similar to ketamine, is neuroprotective in animal models of stroke, trauma, and Parkinsonism, but can induce psychotic behavior and neuronal degeneration; its side effects, likely a result of high affinity for and long dwell time on the NMDAr, preclude its clinical application [15,16]. MK-801 is more potent than ketamine in inhibiting the NMDAr. Because the NMDAr plays a crucial role in mediating excitatory synaptic transmission in the CNS, inhibiting NMDArs profoundly modulates CNS function; MK-801 has been reported to exhibit anticonvulsant and neuroprotective effects during the post-ischemic period, and this pathological effect is believed to be mediated largely by its hypo-glutamatergic/NMDAr-blocking action [17,18]. However, several studies have reported that the psychotic effects of MK-801 are also associated with functional defects in receptors and/or ion channels other than the NMDAr, as well as dysfunction of GABAergic interneurons [19,20]. Parvalbumin-positive GABAergic interneurons are central to shaping communication and controlling excitability within the CNS [21,22].

The Kv3 family of potassium channels is considered important in setting and controlling firing frequency in fast-spiking neurons owing to its fast activation kinetics [23,24]. Kv3.1, one member of this family, is a voltage-gated K^+^ channel involved in the rapid repolarization of the action potential in central neurons; Kv3 channels are required for the fast-spiking phenotype of specific GABAergic interneurons and can carry sufficient repolarizing current to support high-frequency firing. These firing properties contribute to normal gamma oscillations and synchronization of cortical neuronal populations, which are considered signatures of higher-order information processing in the human brain [25–27]. Notably, a decrease of Kv3.1 in the CNS has been reported to be associated with schizophrenia. Kv3.1 knockout mice display increased locomotor activity, stereotypical behavior, and sleep loss, features reminiscent of psychomotor excitement in schizophrenia, and a decrease in Kv3.1 protein is linked to altered fast-spiking parvalbumin-positive interneuron firing patterns and the resulting changes in oscillatory synchrony. Consistent with this, NMDAr antagonists increase spontaneous gamma oscillations in animals, whereas antipsychotic drugs reduce this basal/spontaneous gamma activity, and Kv3.1 knockout mice likewise show increased spontaneous gamma oscillations. Together, this evidence supports the hypothesis that decreased Kv3.1 contributes to the psychotic phenotype observed in schizophrenia [28–30].

In this study, therefore, we examined the effect of MK-801 on Kv3.1 and found that MK-801 directly blocks Kv3.1 with an IC50 of 10.81 μM and a Hill coefficient of 0.89. The block was partly state-dependent, and the steady-state inactivation curve was shifted in the hyperpolarizing direction; recovery from inactivation was also retarded in the presence of MK-801. Given the importance of Kv3.1 in parvalbumin-positive, fast-spiking GABAergic interneurons, these findings suggest that MK-801 may interfere with fast repolarization and alter the firing pattern of these inhibitory interneurons, contributing to the onset and symptoms of schizophrenia.

## Materials and Methods

### Cell culture

Experiments were conducted using Chinese hamster ovary (CHO) cells stably overexpressing Kv3.1. Cells were maintained in Iscove’s modified Dulbecco’s Medium (IMDM) supplemented with 10% fetal bovine serum, HT Supplement (sodium hypoxanthine and thymidine mixture), and G418 disulfate salt solution (each diluted 1:100 in IMDM; 5 mL in 500 mL). Cells were kept at 37°C in a humidified atmosphere of 95% air and 5% CO2. For patch-clamp recordings, cells were treated with 0.05% trypsin-EDTA for 3 min in Ca2+-free normal Tyrode solution (143 mM NaCl, 5.4 mM KCl, 0.33 mM NaH2PO4, 0.5 mM MgCl2, 5 mM HEPES, and 11 mM glucose, pH 7.4 with NaOH), then seeded in a patch-clamp chamber for 30 min before recording.

### Solutions and drugs

For patch-clamp recordings, normal Tyrode solution (143 mM NaCl, 5.4 mM KCl, 0.33 mM NaH2PO4, 0.5 mM MgCl2, 5 mM HEPES, 11 mM glucose, 1.8 mM CaCl2, pH 7.4 with NaOH) was used as the bath solution. (+)-MK-801 hydrogen maleate (M107, Sigma-Aldrich) was dissolved in dimethyl sulfoxide (DMSO, Sigma-Aldrich) to prepare stock solutions (1, 10, and 100 mM), then diluted in the Tyrode bath solution to final concentrations of 1, 3, 10, 30, 100, and 300 µM. The final DMSO concentration after dilution was less than 0.1%, which did not affect Kv3.1 currents. The pipette solution contained 135 mM KCl, 5 mM MgCl2, 10 mM HEPES, 5 mM EGTA, and 5 mM MgATP (pH 7.2 with KOH; osmolality 286 mOsm). All drugs, including the (+)-optical isomer of MK-801, were purchased from Sigma-Aldrich (St. Louis, MO).

### Electrophysiology

Kv3.1 currents were recorded using the conventional whole-cell patch-clamp technique. Membrane voltage was controlled with an EPC-8 amplifier (HEKA Instruments, Germany). Current and voltage signals were digitized using an Axon Digidata 1550B AD/DA converter (Axon Instruments, USA) and stored on a computer. Data were acquired at a sampling rate of 10 kHz after low-pass filtering at 1 kHz. Voltage-clamp control, data acquisition, and analysis were performed using pCLAMP 11 software (Axon Instruments, USA). Patch pipettes were fabricated from Clark borosilicate thin-wall capillaries (Harvard Apparatus Inc., Massachusetts, USA) using a PP-100 puller (Narishige, Japan), yielding a pipette resistance of 1-2 MΩ when filled with internal solution. Kv3.1 currents were elicited by step depolarizations from a holding potential of typically -70 mV; the detailed voltage protocols are shown as insets in each figure.

### Data analysis

Data were analyzed using OriginPro 2016 (OriginLab Corp., Northampton, MA, USA). Results are expressed as means ± SEM. Paired or unpaired Student’s t-tests were used as appropriate to assess statistical significance; p < 0.05 was considered significant.

## Results

### Voltage-dependent inhibition of Kv3.1 currents by MK-801

Kv3.1 currents were elicited by a series of depolarizing step pulses from a holding potential of - 70 mV, followed by repolarization to -35 mV (100 ms) before returning to -70 mV to examine tail currents. Representative recordings in the absence and presence of MK-801 (30 µM) are shown in Figure 1A. MK-801 markedly decreased Kv3.1 currents, particularly at depolarized potentials (≥ 0 mV), with the degree of inhibition increasing with depolarization. Current-voltage (I-V) relationships in the absence and presence of MK-801 are summarized in Figure 1B, shown as current density at the peak (left) and at the end of the 300 ms depolarizing pulse (right). To quantitatively analyze the voltage-dependent inhibition, relative currents (in the presence of MK-801 divided by control) are plotted against membrane voltage in Figure 2. These results indicate that MK-801 blocks Kv3.1 with a steep voltage dependence, particularly above 0 mV.

**Figure 1.**
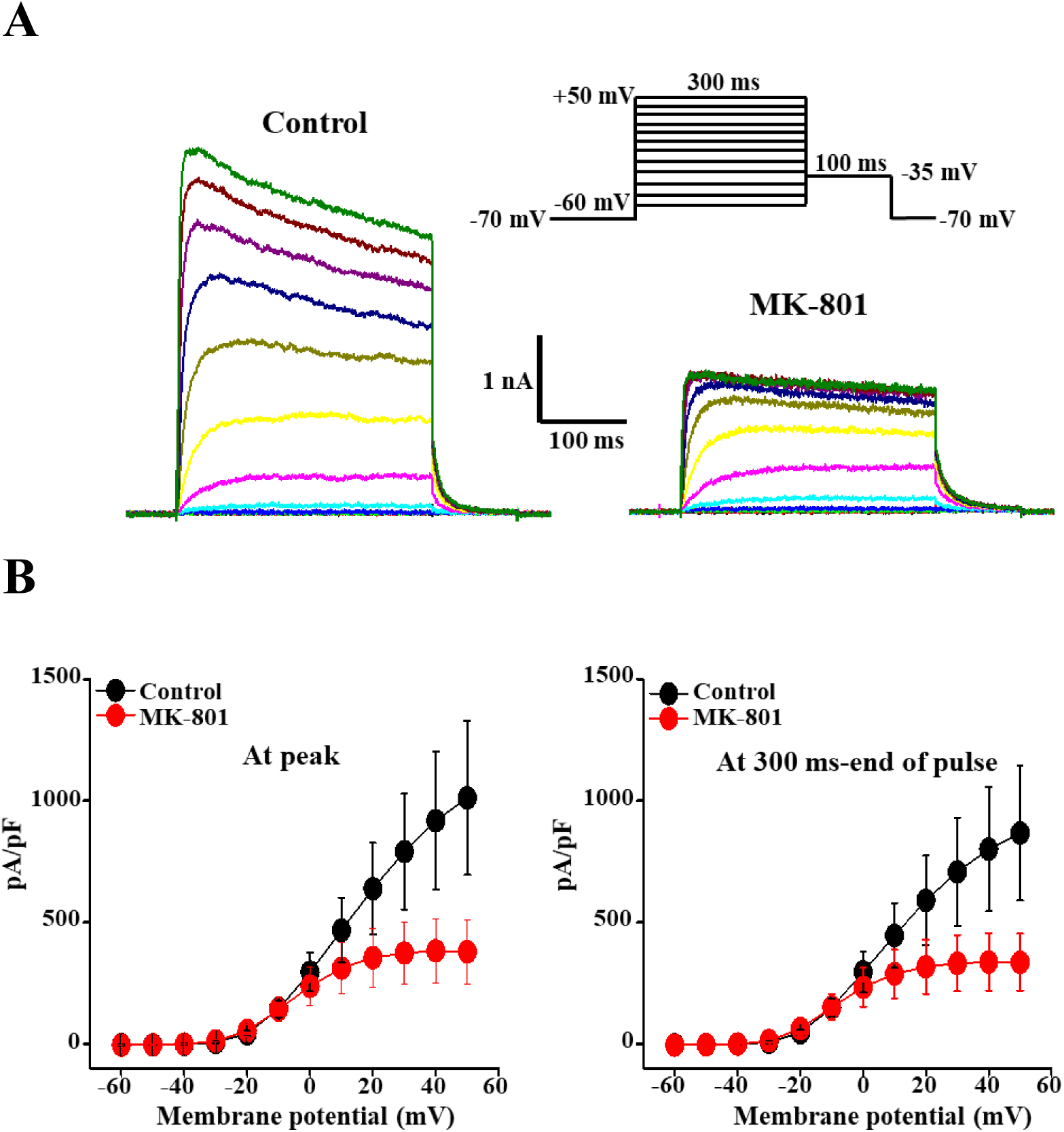
Inhibitory effects of MK-801 on Kv3.1 currents. (A) Superimposed current traces elicited by 300 ms depolarizing pulses between -60 and +50 mV from a holding potential of -70 mV, in the absence and presence of 30 µM MK-801. (B) Current-voltage (I-V) relationships of the peak and quasi-steady-state (end of the 300 ms pulse) Kv3.1 currents in the absence and presence of MK-801 (30 µM). n = 5.

**Figure 2.**
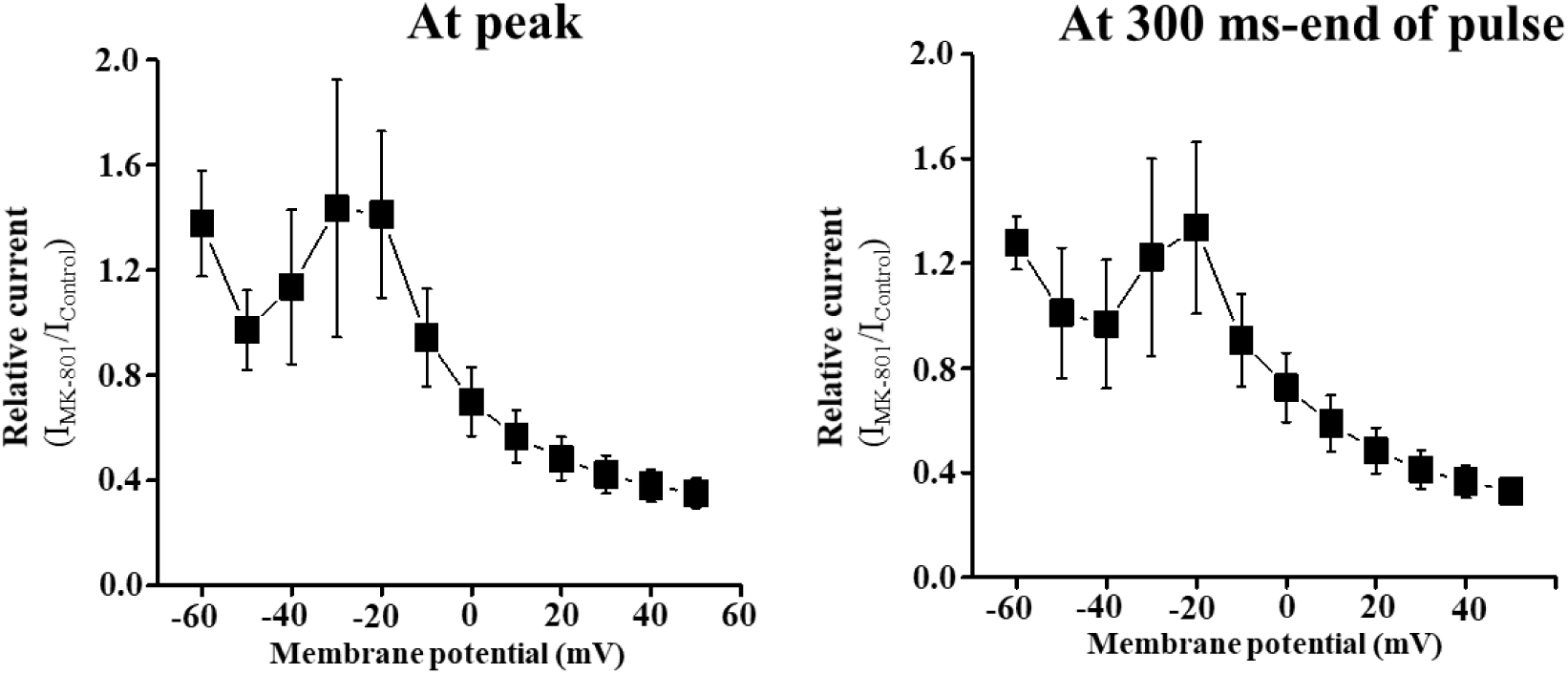
Voltage-dependent inhibition of Kv3.1 currents by MK-801. The degree of inhibition by 30 µM MK-801 is plotted as a function of membrane potential, at peak (left) and quasi-steady-state (right) current levels. n = 5.

### Concentration-response relationship and time course of inhibition

To examine the concentration-response relationship, cumulative concentrations of MK-801 were applied during repetitive (20 s interval) step depolarizations to +50 mV. A representative recording is shown in Figure 3A, and the time course of inhibition by cumulative MK-801 concentrations is shown in Figure 3B (left, peak currents; right, current at the end of the 300 ms pulse). The concentration-inhibition relationship (Figure 3C) yielded an IC50 of 10.81 μM and a Hill coefficient of 0.89, obtained by fitting the data to the Hill equation: ƒ = 1/{1+(Kd/[2])n}, where ƒ is the fraction of inhibition (ƒ = 1-Idrug/Icontrol) at the test potential, Kd is the dissociation constant (IC50), n is the Hill coefficient, and [2] is drug concentration.

**Figure 3.**
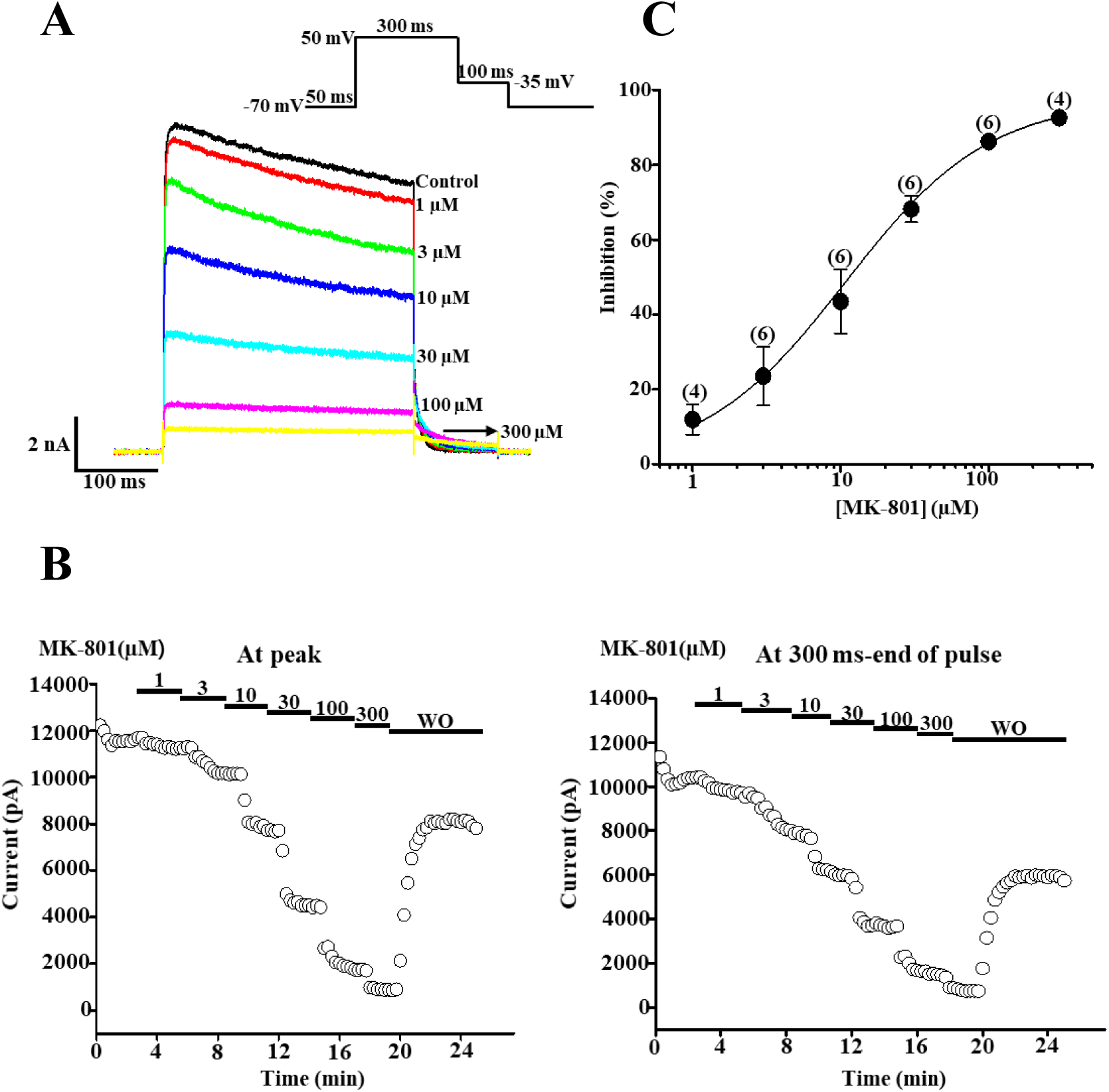
Concentration-response relationship (CRC) and time course of MK-801 inhibition of Kv3.1 currents. (A) Representative recordings showing the effect of cumulative MK-801 concentrations (1-300 µM), elicited by 300 ms depolarizing pulses from -70 to +50 mV at 20 s intervals. (B) Time course of inhibition and washout recovery (peak and quasi-steady-state values). (C) Concentration-dependence of inhibition; numbers in parentheses indicate the number of cells tested. Data were fit to the Hill equation, yielding an IC50 of 10.81 µM and a Hill coefficient of 0.89. n = 6.

### Effects on activation and deactivation

The voltage dependence of steady-state activation was examined using a double-pulse protocol (Figure 4A inset). Tail current amplitudes after various test potentials were normalized to obtain steady-state activation curves in the absence and presence of MK-801 (30 µM) (Figure 4B).

**Figure 4.**
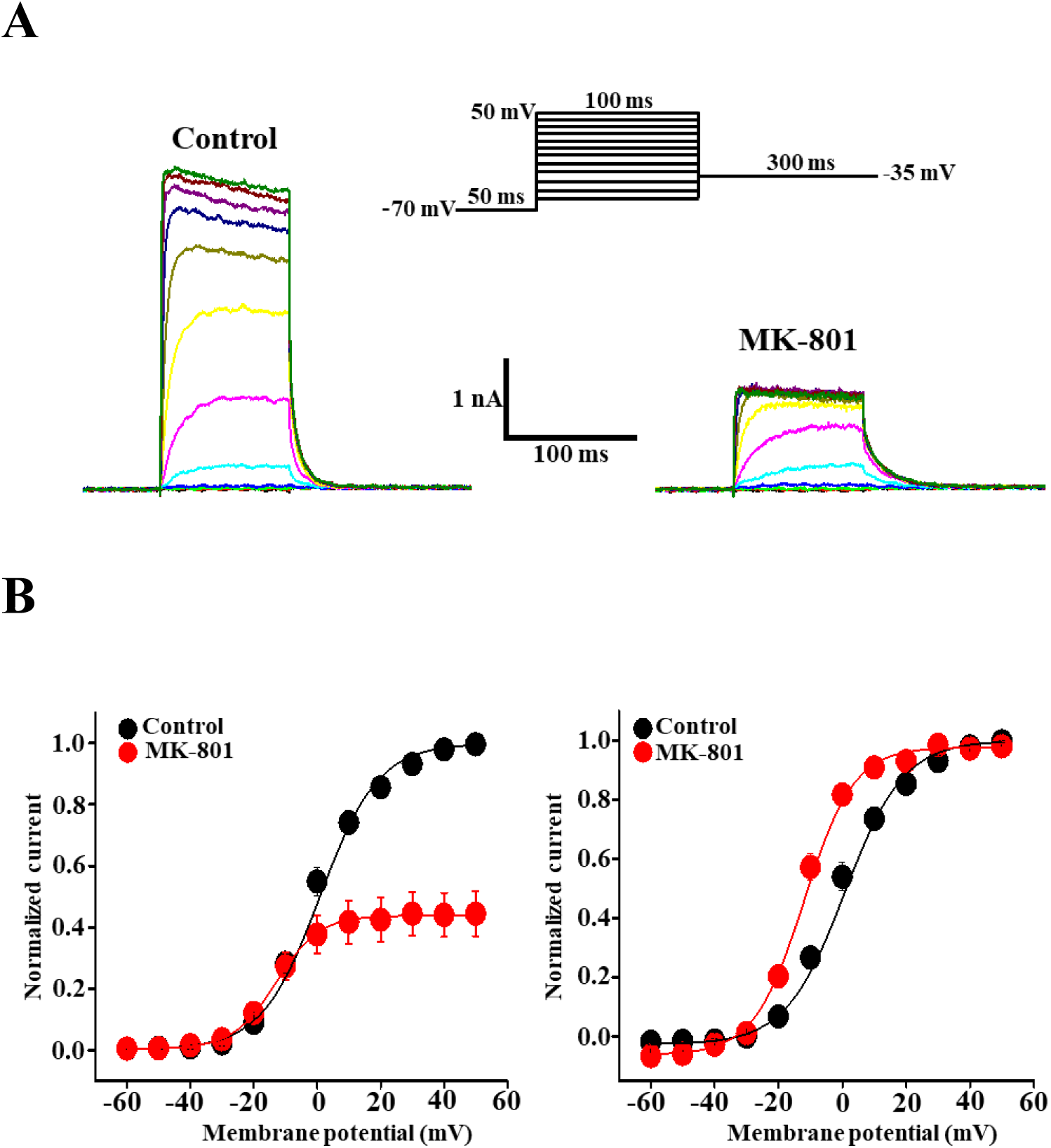
Effects of MK-801 on steady-state activation of Kv3.1. (A) Representative current traces elicited by a double-pulse protocol (inset) under control conditions and in the presence of 30 µM MK-801. (B) Normalized current amplitude (an estimate of channel activation) plotted against test pulse potential; smooth lines are Boltzmann fits: y = A2+(A1-A2)/(1+exp((x-x0)/dx)). Left panel: currents normalized to peak tail current at +50 mV in the absence of MK-801. Right panel: currents normalized to peak tail current of the corresponding condition. n = 5.

Because MK-801 inhibited Kv3.1 only at strongly depolarized potentials (≥ 0 mV), the apparent activation curve shifted leftward (Figure 4B, right panel); however, we interpret this as a consequence of the steep voltage dependence of MK-801 block rather than a true activating shift. The time courses of activation and deactivation were also compared. Under control conditions, the activation time constant was voltage-dependent, becoming faster with depolarization (Figure 5), and MK-801 did not affect activation kinetics. In contrast, the deactivation time course during repolarization to -35 mV after a +50 mV depolarization was significantly slower in the presence of MK-801 (Figure 6).

**Figure 5.**
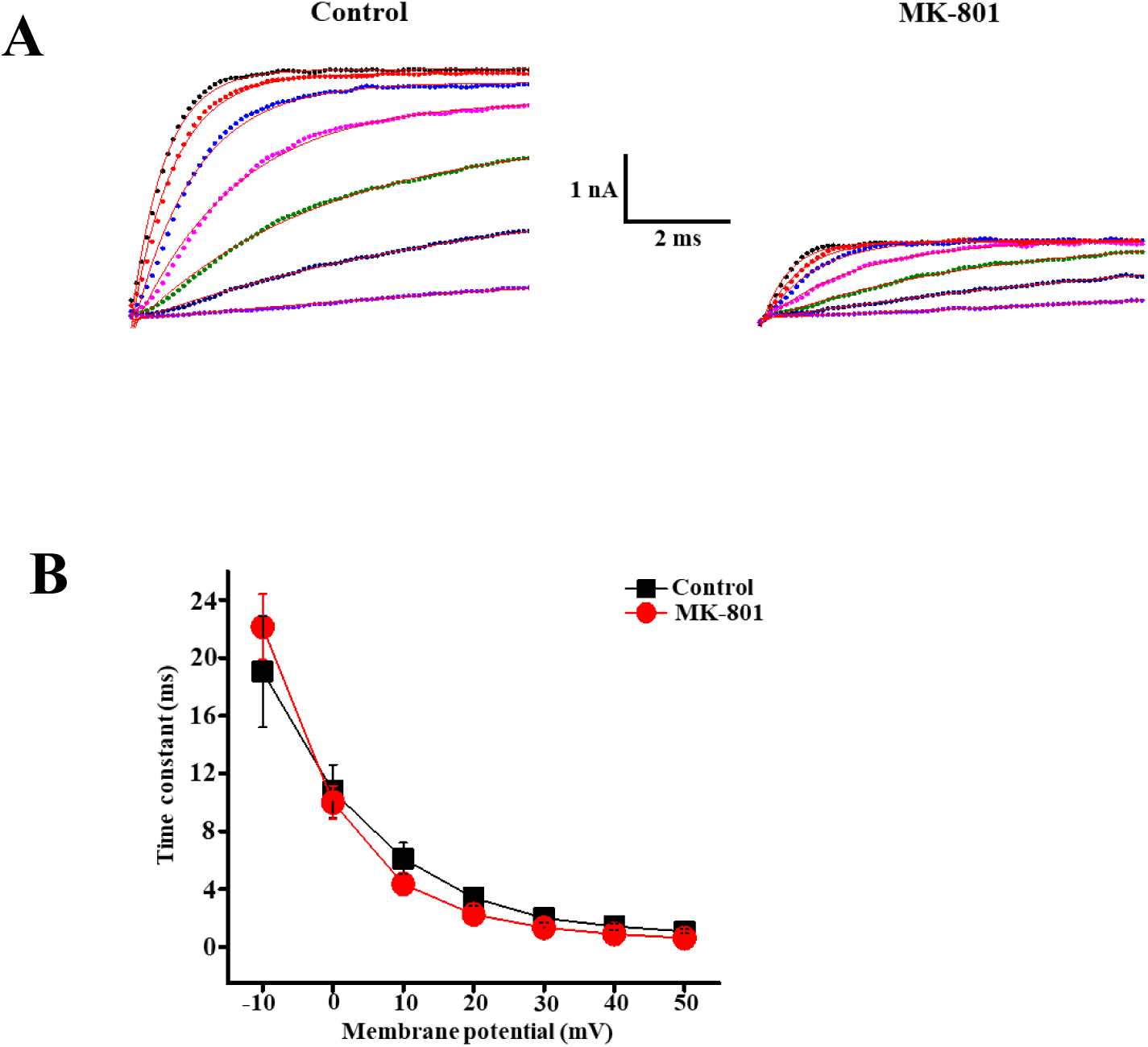
Effects of MK-801 on activation time courses of Kv3.1. (A) Superimposed current traces elicited by depolarizing pulses (-10 to +50 mV) from a holding potential of -70 mV, in the absence (left) and presence (right) of MK-801; smooth lines are single-exponential fits. (B) Activation time constants across membrane potentials, with and without MK-801. n = 5.

**Figure 6.**
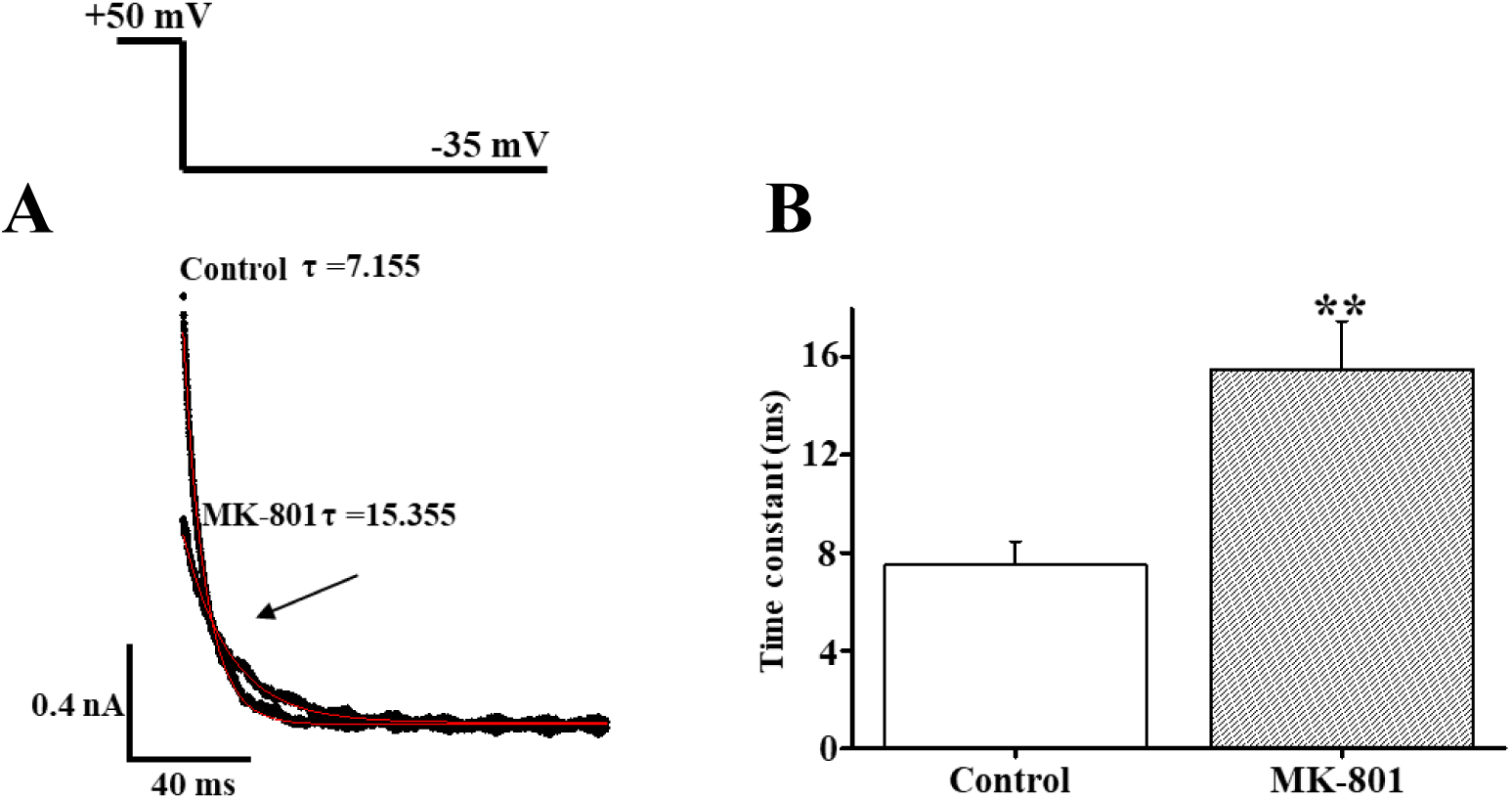
Effects of MK-801 on the deactivation time course of Kv3.1. (A) Representative deactivating tail currents during repolarization to -35 mV following depolarization to +50 mV, in the absence and presence of MK-801, with single-exponential fits. (B) Deactivation time constants; arrow indicates cross-over of tail currents due to the retarded deactivation time course with MK-801. n = 5. **P < 0.01 (paired t-test).

### Effects on inactivation of Kv3.1

Steady-state inactivation was examined using a double-pulse protocol in which 10 s test pulses of varying voltage preceded a brief depolarizing pulse (+40 mV) (Figure 7A inset). MK-801 shifted the steady-state inactivation curve leftward (Figure 7B); the voltage for half-inactivation was -25.49 ± 1.22 mV under control conditions and -40.17 ± 1.58 mV in the presence of MK-801. In contrast, the time course of inactivation, obtained by single-exponential fitting, was minimally affected by MK-801 (Figure 8): the inactivation time constant at +20 mV was 2133.98 ± 363.39 ms under control conditions and 2238.12 ± 418.02 ms with MK-801.

**Figure 7.**
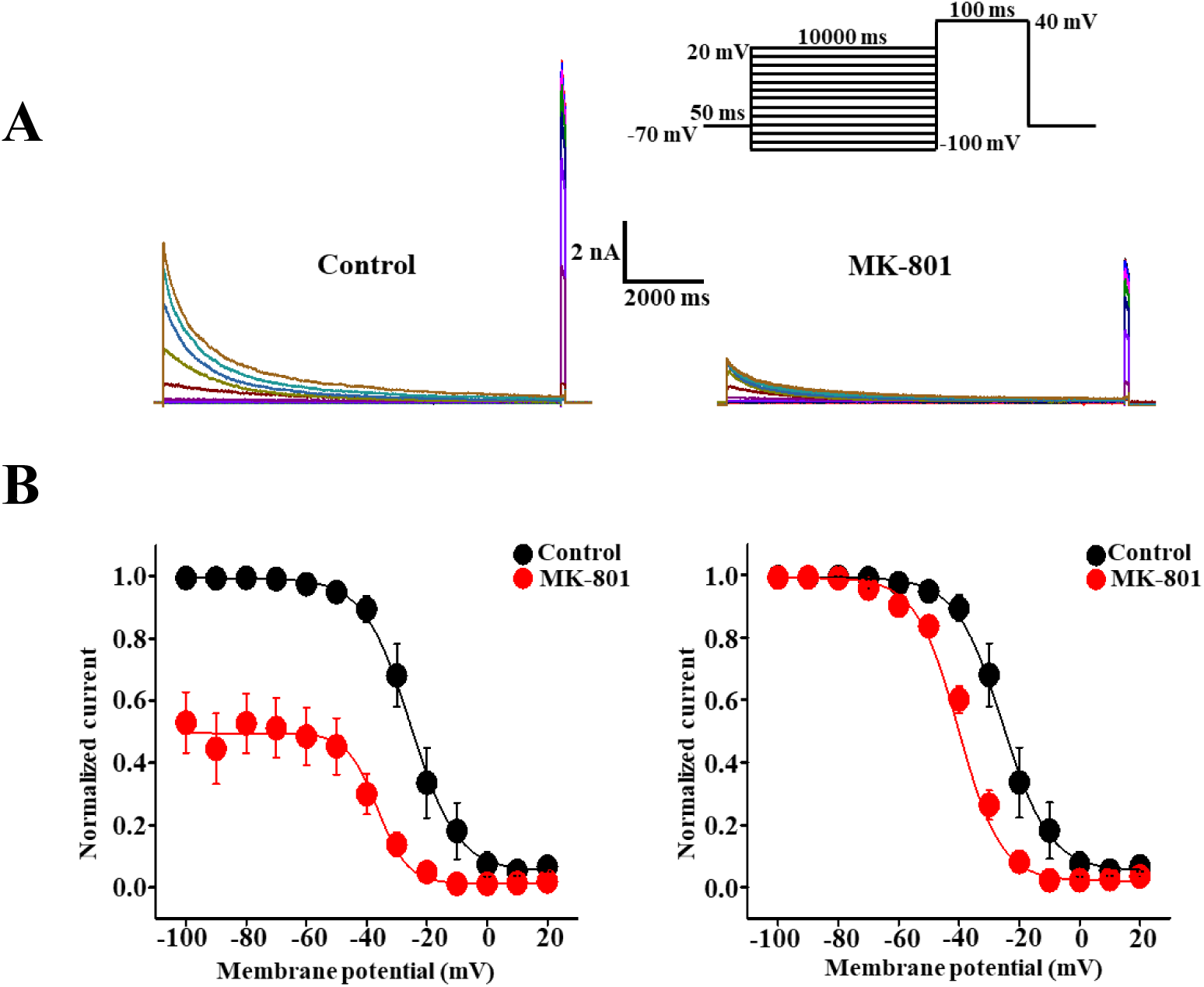
Effects of MK-801 on steady-state inactivation of Kv3.1. (A) Superimposed current traces from a double-pulse protocol (inset) for measuring steady-state inactivation, in the absence (left) and presence (right) of MK-801 (30 µM). (B) Normalized current (an estimate of steady-state inactivation) plotted against pre-pulse potential; smooth lines are Boltzmann fits (as in Figure 4). Left panel normalized to peak current without MK-801; right panel normalized to peak current of the corresponding condition. Note the leftward shift with MK-801. n = 6.

**Figure 8.**
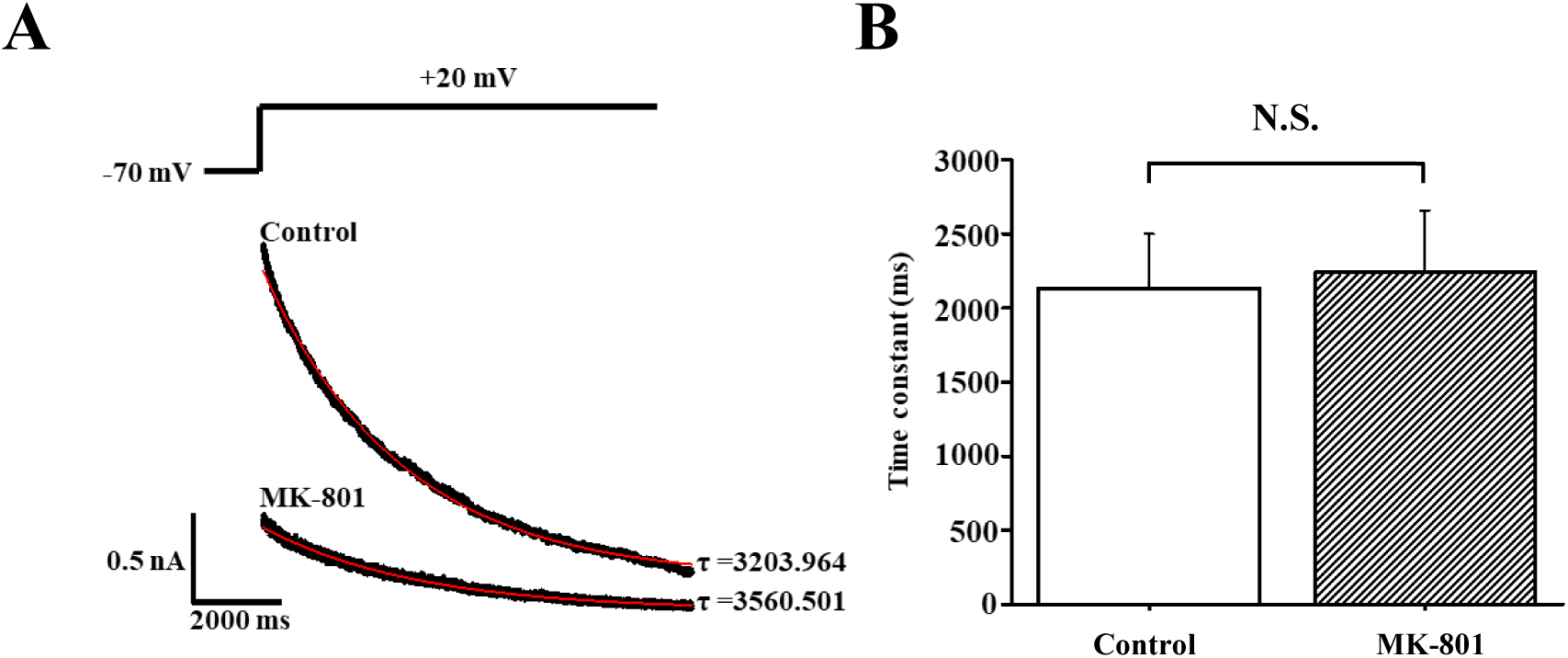
Effects of MK-801 on the inactivation time course of Kv3.1. (A) Representative current traces showing the slow inactivation time course during a long depolarizing step to +20 mV, in the absence and presence of MK-801, with single-exponential fits. (B) Inactivation time constants at +20 mV, with and without MK-801. n = 4.

### Use-dependent block of Kv3.1 by MK-801

To determine whether the blocking effect of MK-801 is use- or state-dependent, trains of depolarizing pulses (20 pulses at 1 or 2 Hz) were applied in the absence and presence of MK-801 (Figure 9A, B). After MK-801 application, the first depolarizing pulse was applied 3 min later to allow sufficient interaction time; approximately 50% inhibition was already present at the first pulse. In the presence of MK-801, Kv3.1 currents decreased progressively over the pulse train (Figure 9A-C), suggesting that part of the MK-801 block of Kv3.1 is use-dependent, while another part is use- or state-independent.

**Figure 9.**
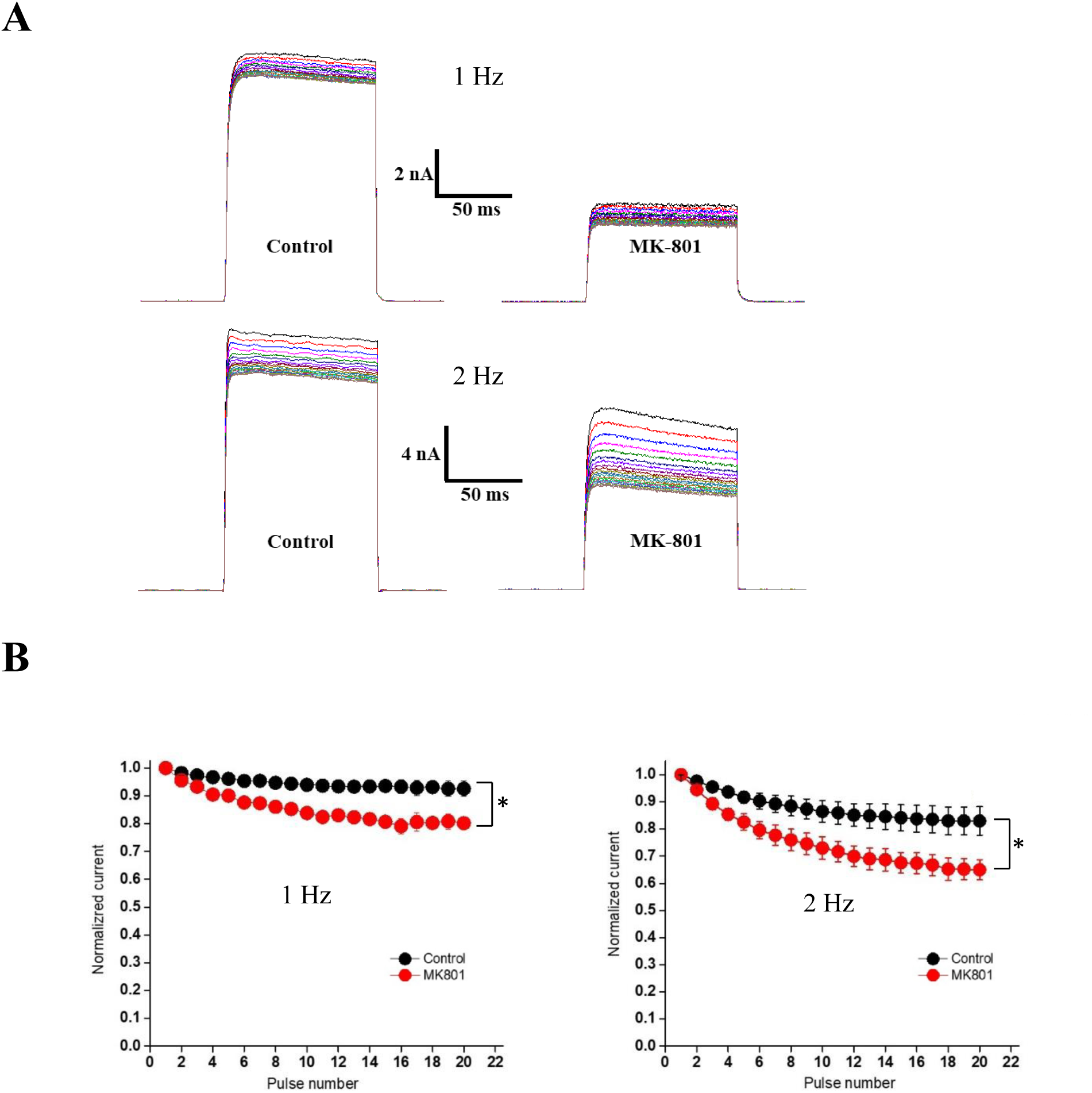
Use-dependence of MK-801 inhibition of Kv3.1 currents. Twenty repetitive 100 ms depolarizing pulses to +40 mV from a holding potential of -70 mV were applied at 1 Hz (A) and 2 Hz (B) with and without 30 µM MK-801. (A, B) Representative superimposed current traces. (C) Peak amplitudes at each pulse train frequency, normalized to the first pulse, plotted against pulse number. n = 3. *P < 0.05 (paired t-test).

### Kinetics of recovery from inactivation

To further test whether the use/state-dependent component of block reflects preferential interaction of MK-801 with the inactivated state of Kv3.1, recovery from inactivation was examined using a long (10 s) conditioning pulse followed by test pulses at varying intervals (Figure 10A inset). Because recovery followed a two-component (fast and slow) time course, 50 ms and 1 s interval protocols were used to resolve the fast and slow components, respectively.

**Figure 10.**
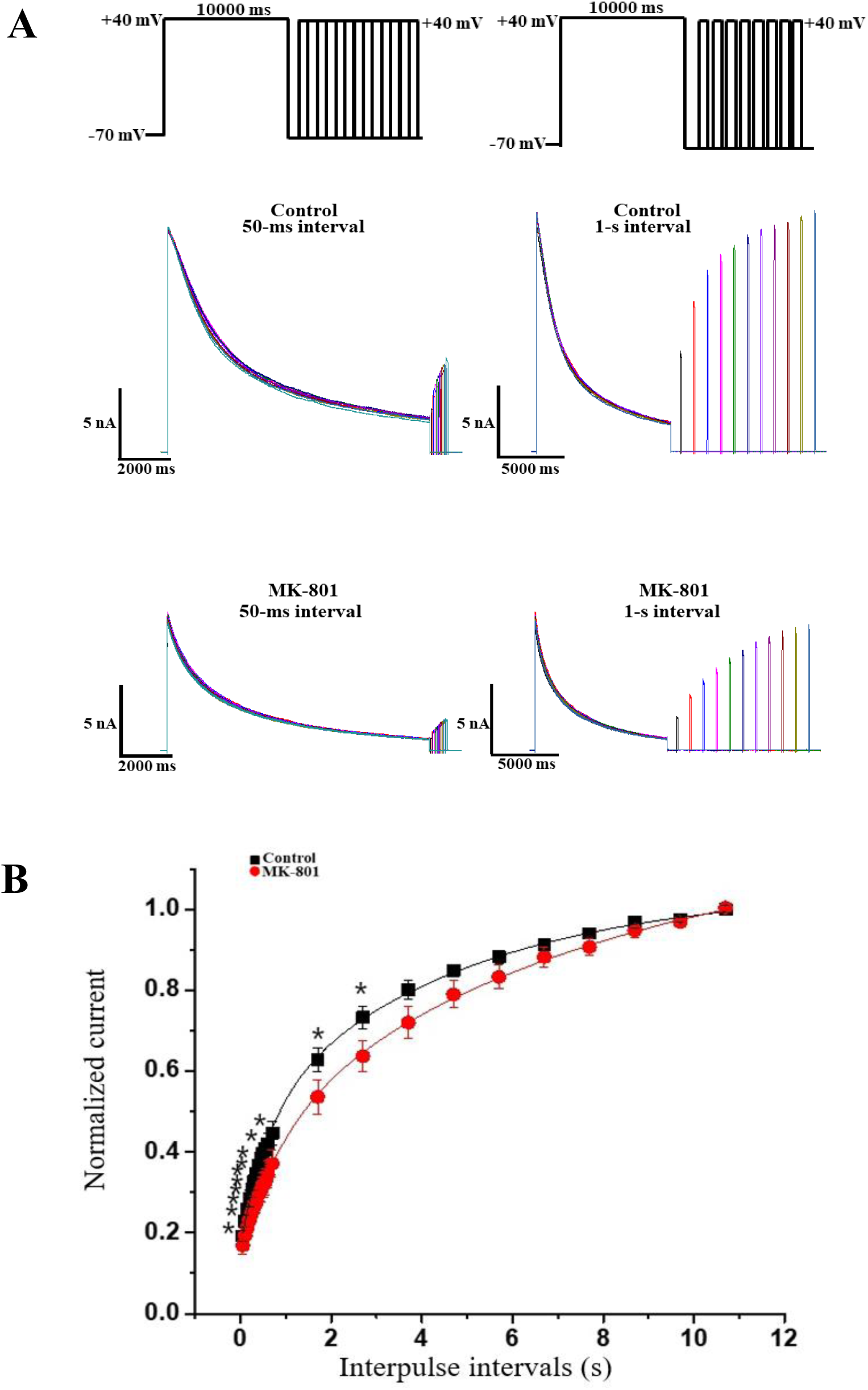
Effects of MK-801 on the time course of recovery from inactivation of Kv3.1. Recovery was measured using a double-pulse protocol (inset): a 10 s depolarizing pre-pulse to +40 mV from a holding potential of -70 mV was followed by a second pulse after variable intervals (50 ms-11 s). (A) Representative current tracings showing fast (left) and slow (right) components of recovery, in the absence (upper) and presence (lower) of MK-801. (B) Peak currents from the second test pulse, normalized to the first pre-pulse and plotted against inter-pulse interval; smooth lines are double-exponential fits. n = 5. *p < 0.05. (paired t-test)

MK-801 clearly retarded the recovery time course (Figure 10A, B). Recovery time constants were 0.64 ± 0.079 s (fast) and 4.71 ± 0.49 s (slow) under control conditions, versus 0.95 ± 0.13 s (fast) and 7.77 ± 1.29 s (slow) in the presence of MK-801. These results suggest that retarded recovery from inactivation contributes to the use/state-dependent inhibition of Kv3.1 by MK-801.

## Discussion

PCP and its non-competitive NMDAr antagonist derivatives, including MK-801 and ketamine, induce positive and negative symptoms in healthy individuals resembling schizophrenia and successfully generate an animal model of the disease [8–10]. This has led to considerable interest in the glutamate (hypo-glutamatergic) hypothesis of schizophrenia [6,7]. However, growing evidence indicates that "off-target" actions of NMDAr antagonists also contribute to the pathogenesis of schizophrenia; for example, serotonin 5-HT2 and dopamine D2 receptors have been reported to be activated by these antagonists [3,9]. Here we identify another candidate off-target, Kv3.1, which is richly expressed in parvalbumin-positive, fast-spiking GABAergic interneurons that are concentrated in the prefrontal cortex and implicated in higher cognitive and social functions [21–27]. Impaired Kv3.1 function has been reported to be closely associated with the onset of schizophrenia symptoms [28–31]. In this study, we demonstrate voltage- and state (use)-dependent inhibition of Kv3.1 by MK-801 at pharmacologically relevant concentrations, suggesting that Kv3.1 may represent an additional target contributing to the pathogenesis of schizophrenia induced by PCP-derivative NMDAr antagonists.

### MK-801 directly blocks Kv3.1 in a voltage- and use (state)-dependent manner at pharmacologically relevant concentrations

Because CHO cells do not express the NMDAr, the effects of MK-801 on Kv3.1 currents reported here are not mediated by the NMDAr but reflect a direct interaction. The rapid inhibition time course (Figure 3B) further supports this direct interaction. Inhibition was concentration-dependent with an IC50 of 10.81 µM and a Hill coefficient of 0.89 (Figure 3), a potency substantially higher than that reported for MK-801 on Kv1.5 overexpressed in CHO cells or on native Kv currents of rat mesenteric arterial myocytes (157 and 89 µM, respectively) [2,14] . In MK-801-induced animal models of schizophrenia, doses of approximately 0.5 mg/kg have been used [13]. At a lower dose (0.1 mg/kg) in mice, blood plasma concentrations of MK-801 have been reported to be 7.4 ± 1.6 and 6.8 ± 1.9 ng/mL after subcutaneous (s.c.) and intraperitoneal (i.p.) administration, respectively, with correspondingly higher brain tissue concentrations (57.6 ± 7.6 and 37.9 ± 8.9 ng/mL, respectively; 57.6 ng/mL corresponds to 0.26 µM) [32,33]. These data suggest that brain concentrations of MK-801 would exceed 1.3 µM at a dose of 0.5 mg/kg, indicating that the blockade of Kv3.1 reported here occurs at pharmacologically relevant concentrations with respect to MK-801-induced models of schizophrenia.

The blockade of Kv3.1 by MK-801 was rapidly reversible and concentration-dependent (Figure 3). Given the fast kinetics of block and washout, and the absence of functional NMDArs in CHO cells, we interpret this as a direct interaction between MK-801 and Kv3.1. The block was also steeply voltage-dependent (Figures 1, 2), suggesting that MK-801 senses the transmembrane potential; MK-801 preferentially suppressed Kv3.1 at potentials ≥ 0 mV (Figure 1). MK-801 did not appear to block the open state or inhibit activation, as it neither accelerated relaxation or inactivation (Figures 1, 3, 8), shifted the steady-state activation curve rightward (Figure 4), nor slowed the activation time course (Figure 5). Instead, MK-801 appears to interact with an extracellular site on Kv3.1 associated with facilitation or stabilization of the inactivated state, consistent with the leftward shift of the steady-state inactivation curve (Figure 7) and the retarded recovery from inactivation (Figure 10). These findings also help explain the use- or state-dependent inhibition shown in Figure 9. MK-801 additionally retarded the deactivation time course, producing a "cross-over" of tail currents characteristic of open-channel block. We propose that MK-801 interacts with an extracellular site near, but not within, the channel pore, facilitating a voltage-dependent conformational transition to a non-conducting, inactivated-like state; upon repolarization, prior dissociation of MK-801 may be required before Kv3.1 can return from the activated to the closed state (deactivation). This model remains speculative and warrants further study.

### Neurotransmitters in the pathophysiology of schizophrenia

The NMDA receptor plays a well-established role in mediating excitatory signaling in the CNS, and MK-801 has been widely used to generate schizophrenia models via NMDA receptor antagonism, and its behavioral and cognitive effects have been extensively characterized [34].

However, the pathophysiology of schizophrenia is not limited to NMDA receptor dysfunction; dopamine, serotonin, and glutamic acid, as well as aspartic acid, glycine, and GABA, have all been implicated in the associated neurochemical imbalance. For example, positive symptoms such as hallucinations and delusions have been linked to increased subcortical dopamine release and D2 receptor activation [35], whereas negative symptoms such as anhedonia, reduced motivation, and impaired language have been linked to decreased D1 receptor activation in the prefrontal cortex, with D3 receptor alterations also implicated. Dopamine agonists increase D2-high receptors [36], a change considered a basic requirement for the development of dopamine-hypersensitivity psychosis, and dysregulated dopamine signaling in the basal ganglia is considered an important intrinsic feature of schizophrenia pathology. Ketamine and PCP have also been reported to produce broader, non-specific neurochemical perturbations involving the glutamatergic, dopaminergic, and serotonergic systems [9,37,38].

### Cortical GABA neurons and gamma oscillations: role of Kv3.1

Activity of parvalbumin-containing GABA neurons is closely linked to the generation of cortical network oscillations [39]. Gamma-band oscillations (30-80 Hz) are considered essential for cortical information transmission and processing, and disruption of this synchronization in schizophrenia is thought to contribute to cognitive impairment [40]. As introduced above, Kv3.1 is essential for the normal regulation of parvalbumin-containing GABA neurons, and its reduction in the CNS has been linked to schizophrenia-like phenotypes. Notably, the behavioral excess observed in Kv3.1 knockout mice parallels the elevation of spontaneous gamma-band oscillations induced by NMDA receptor antagonists, an excess that antipsychotic drugs normalize [41], and Kv3.1 knockout mice likewise show increased spontaneous gamma oscillations [30]. Together, this evidence supports the hypothesis that decreased Kv3.1 contributes to the psychotic phenotype observed in schizophrenia.

## Conclusion

Taken together, these results indicate that the PCP-derivative NMDAr antagonist MK-801 blocks Kv3.1 in a concentration-, voltage-, and state-dependent manner at clinically relevant concentrations. Given the regional expression of Kv3.1, its contribution to fast-spiking cortical GABAergic interneuron firing and cortical gamma oscillations, and the role of these processes in higher cortical functions, impaired Kv3.1 function represents a plausible mechanism contributing to the schizophrenia phenotype. We therefore propose Kv3.1 as a relevant target in the pathogenesis of schizophrenia and a candidate for future therapeutic development, consistent with recent structural and clinical evidence supporting Kv3 channel modulators as candidate schizophrenia therapeutics [42–44].

## Author Contributions

Conceptualization: Tae Jun Park, Sang Woong Park, Young Min Bae Methodology: Tae Jun Park, Sang Woong Park, Mi Seon Seo, Jin Ryeol An

Investigation: Tae Jun Park, Sang Woong Park, Mi Seon Seo, Jin Ryeol An, Solah Park, Yeji Lim, Hyeryeong Lee, Jeongbin Kim, Harim Kim

Data Curation: Tae Jun Park, Sang Woong Park Formal Analysis: Tae Jun Park, Sang Woong Park

Resources: Jin O-Uchi, Dong Hyun Kim, Bok Hee Choi, Young Min Bae Writing – Original Draft: Tae Jun Park, Sang Woong Park

Writing – Review & Editing: Tae Jun Park, Sang Woong Park, Bok Hee Choi, Jin O-Uchi, Dong Hyun Kim, Young Min Bae

## References

[1] R. Brisch, A. Saniotis, R. Wolf, H. Bielau, H.G. Bernstein, J. Steiner, et al. The role of dopamine in schizophrenia from a neurobiological and evolutionary perspective: old fashioned, but still in vogue. Front Psychiatry 5 (2014) 47.

[2] H. Lin, J.G. Kim, S.W. Park, H.J. Noh, J.M. Kim, C.Y. Yoon, et al. Enhancement of 5-HT(2A) receptor function and blockade of Kv1.5 by MK801 and ketamine: implications for PCP derivative-induced disease models. Exp Mol Med 50 (2018) 1–8.

[3] R.A. McCutcheon, J.H. Krystal and O.D. Howes. Dopamine and glutamate in schizophrenia: biology, symptoms and treatment. World Psychiatry 19 (2020) 15–33.

[4] P.W. Chiu, S.S.Y. Lui, K.S.Y. Hung, R.C.K. Chan, Q. Chan, P.C. Sham, et al. In vivo gamma-aminobutyric acid and glutamate levels in people with first-episode schizophrenia: A proton magnetic resonance spectroscopy study. Schizophr Res 193 (2018) 295–303.

[5] A.C. Yang and S.J. Tsai. New Targets for Schizophrenia Treatment beyond the Dopamine Hypothesis. Int J Mol Sci 18 (2017)

[6] W. Hu, M.L. MacDonald, D.E. Elswick and R.A. Sweet. The glutamate hypothesis of schizophrenia: evidence from human brain tissue studies. Ann N Y Acad Sci 1338 (2015) 38–57.

[7] J.T. Kantrowitz and D.C. Javitt. N-methyl-d-aspartate (NMDA) receptor dysfunction or dysregulation: the final common pathway on the road to schizophrenia? Brain Res Bull 83 (2010) 108–121.

[8] C. Bondi, M. Matthews and B. Moghaddam. Glutamatergic animal models of schizophrenia. Curr Pharm Des 18 (2012) 1593–1604.

[9] S. Kapur and P. Seeman. NMDA receptor antagonists ketamine and PCP have direct effects on the dopamine D(2) and serotonin 5-HT(2)receptors-implications for models of schizophrenia. Mol Psychiatry 7 (2002) 837–844.

[10] D. Cadinu, B. Grayson, G. Podda, M.K. Harte, N. Doostdar and J.C. Neill. NMDA receptor antagonist rodent models for cognition in schizophrenia and identification of novel drug treatments, an update. Neuropharmacology 142 (2018) 41–62.

[11] J.A. Pratt, C. Winchester, A. Egerton, S.M. Cochran and B.J. Morris. Modelling prefrontal cortex deficits in schizophrenia: implications for treatment. Br J Pharmacol 153 Suppl 1 (2008) S465–470.

[12] J.W. Newcomer, N.B. Farber and J.W. Olney. NMDA receptor function, memory, and brain aging. Dialogues Clin Neurosci 2 (2000) 219–232.

[13] E.M. Eyjolfsson, E. Brenner, D. Kondziella and U. Sonnewald. Repeated injection of MK801: an animal model of schizophrenia? Neurochem Int 48 (2006) 541–546.

[14] J.M. Kim, S.W. Park, H.Y. Lin, K.C. Shin, D.J. Sung, J.G. Kim, et al. Blockade of voltage-gated K(+) currents in rat mesenteric arterial smooth muscle cells by MK801. J Pharmacol Sci 127 (2015) 92–102.

[15] C. Seillier, F. Lesept, O. Toutirais, F. Potzeha, M. Blanc and D. Vivien. Targeting NMDA Receptors at the Neurovascular Unit: Past and Future Treatments for Central Nervous System Diseases. Int J Mol Sci 23 (2022)

[16] P. Kovacic and R. Somanathan. Clinical physiology and mechanism of dizocilpine (MK-801): electron transfer, radicals, redox metabolites and bioactivity. Oxid Med Cell Longev 3 (2010) 13–22.

[17] J. Liu, L. Chang, Y. Song, H. Li and Y. Wu. The Role of NMDA Receptors in Alzheimer’s Disease. Front Neurosci 13 (2019) 43.

[18] G. Lee and Y. Zhou. NMDAR Hypofunction Animal Models of Schizophrenia. Front Mol Neurosci 12 (2019) 185.

[19] J.T. Li, Y.A. Su, H.L. Wang, Y.Y. Zhao, X.M. Liao, X.D. Wang, et al. Repeated Blockade of NMDA Receptors During Adolescence Impairs Reversal Learning and Disrupts GABAergic Interneurons in Rat Medial Prefrontal Cortex. Front Mol Neurosci 9 (2016) 17.

[20] S.M. Cohen, R.W. Tsien, D.C. Goff and M.M. Halassa. The impact of NMDA receptor hypofunction on GABAergic neurons in the pathophysiology of schizophrenia. Schizophr Res 167 (2015) 98–107.

[21] Z.J. Huang and A. Paul. The diversity of GABAergic neurons and neural communication elements. Nat Rev Neurosci 20 (2019) 563–572.

[22] C. Le Magueresse and H. Monyer. GABAergic interneurons shape the functional maturation of the cortex. Neuron 77 (2013) 388–405.

[23] B. Rudy and C.J. McBain. Kv3 channels: voltage-gated K+ channels designed for high-frequency repetitive firing. Trends Neurosci 24 (2001) 517–526.

[24] A. Erisir, D. Lau, B. Rudy and C.S. Leonard. Function of specific K(+) channels in sustained high-frequency firing of fast-spiking neocortical interneurons. J Neurophysiol 82 (1999) 2476–2489.

[25] L.K. Kaczmarek and Y. Zhang. Kv3 Channels: Enablers of Rapid Firing, Neurotransmitter Release, and Neuronal Endurance. Physiol Rev 97 (2017) 1431–1468.

[26] H.M. Lee, O.H. Chai, S.J. Hahn and B.H. Choi. Antidepressant drug paroxetine blocks the open pore of Kv3.1 potassium channel. Korean J Physiol Pharmacol 22 (2018) 71–80.

[27] K. Boddum, C. Hougaard, J. Xiao-Ying Lin, N.L. von Schoubye, H.S. Jensen, M. Grunnet, et al. K(v)3.1/K(v)3.2 channel positive modulators enable faster activating kinetics and increase firing frequency in fast-spiking GABAergic interneurons. Neuropharmacology 118 (2017) 102–112.

[28] M. Olszewski, W. Dolowa, P. Matulewicz, S. Kasicki and M.J. Hunt. NMDA receptor antagonist-enhanced high frequency oscillations: are they generated broadly or regionally specific? Eur Neuropsychopharmacol 23 (2013) 1795–1805.

[29] R.H. Joho, C.S. Ho and G.A. Marks. Increased gamma- and decreased delta-oscillations in a mouse deficient for a potassium channel expressed in fast-spiking interneurons. J Neurophysiol 82 (1999) 1855–1864.

[30] M. Yanagi, R.H. Joho, S.A. Southcott, A.A. Shukla, S. Ghose and C.A. Tamminga. Kv3.1-containing K(+) channels are reduced in untreated schizophrenia and normalized with antipsychotic drugs. Mol Psychiatry 19 (2014) 573–579.

[31] O. Marin. Parvalbumin interneuron deficits in schizophrenia. Eur Neuropsychopharmacol 82 (2024) 44–52.

[32] B. Svalbe, G. Stelfa, E. Vavers, B. Zvejniece, S. Grinberga, E. Sevostjanovs, et al. Effects of the N-methyl-d-aspartate receptor antagonist, MK-801, on spatial memory and influence of the route of administration. Behav Brain Res 372 (2019) 112067.

[33] B. Svalbe, S. Grinberga, G. Stelfa, E. Vavers, B. Zvejniece, E. Sevostjanovs, et al. Data on analysis of MK-801 bioavailability in mouse plasma and brain tissue by ultra-performance liquid chromatography-tandem mass spectrometry. Data Brief 27 (2019) 104623.

[34] A. Janus, K. Lustyk and K. Pytka. MK-801 and cognitive functions: Investigating the behavioral effects of a non-competitive NMDA receptor antagonist. Psychopharmacology (Berl) 240 (2023) 2435–2457.

[35] Y.Q. Zhang, W.P. Lin, L.P. Huang, B. Zhao, C.C. Zhang and D.M. Yin. Dopamine D2 receptor regulates cortical synaptic pruning in rodents. Nat Commun 12 (2021) 6444.

36. G. Collo, A. Mucci, G.M. Giordano, E. Merlo Pich and S. Galderisi. Corrigendum: Negative Symptoms of Schizophrenia and Dopaminergic Transmission: Translational Models and Perspectives Opened by iPSC Techniques. Front Neurosci 15 (2021) 729082.

[37] N.C. Andreasen, P. Nopoulos, S. Schultz, D. Miller, S. Gupta, V. Swayze, et al. Positive and negative symptoms of schizophrenia: past, present, and future. Acta Psychiatr Scand Suppl 384 (1994) 51–59.

[38] Y. Okubo, T. Suhara, K. Suzuki, K. Kobayashi, O. Inoue, O. Terasaki, et al. Decreased prefrontal dopamine D1 receptors in schizophrenia revealed by PET. Nature 385 (1997) 634–636.

[39] D.A. Lewis, A.A. Curley, J.R. Glausier and D.W. Volk. Cortical parvalbumin interneurons and cognitive dysfunction in schizophrenia. Trends Neurosci 35 (2012) 57–67.

[40] G. Gonzalez-Burgos, T. Hashimoto and D.A. Lewis. Alterations of cortical GABA neurons and network oscillations in schizophrenia. Curr Psychiatry Rep 12 (2010) 335–344.

[41] M. Olszewski, J. Piasecka, S.A. Goda, S. Kasicki and M.J. Hunt. Antipsychotic compounds differentially modulate high-frequency oscillations in the rat nucleus accumbens: a comparison of first- and second-generation drugs. Int J Neuropsychopharmacol 16 (2013) 1009–1020.

[42] M. Yanagi and M. Hashimoto. Dysfunctional Parvalbumin Neurons in Schizophrenia and the Pathway to the Clinical Application of Kv3 Channel Modulators. Int J Mol Sci 25 (2024)

[43] Y.T. Chen, M.R. Hong, X.J. Zhang, J. Kostas, Y. Li, R.L. Kraus, et al. Identification, structural, and biophysical characterization of a positive modulator of human Kv3.1 channels. Proc Natl Acad Sci U S A 120 (2023) e2220029120.

[44] S.J. Kaar, J.F. Nottage, I. Angelescu, T.R. Marques and O.D. Howes. Gamma Oscillations and Potassium Channel Modulation in Schizophrenia: Targeting GABAergic Dysfunction. Clin EEG Neurosci 55 (2024) 203–213.

